# ‘rbioacc’: an ‘R’-package to analyze toxicokinetic data

**DOI:** 10.1101/2021.09.08.459421

**Authors:** A. Ratier, V. Baudrot, M. Kaag, A. Siberchicot, C. Lopes, S. Charles

## Abstract

1. ‘rbioacc’ is an ‘R’-package dedicated to the analysis of experimental data collected from bioaccumulation tests during which organisms are exposed to a chemical (exposure phase) and then put into a clean media (depuration phase). Internal concentrations are regularly measured over time all along the experiment.
2. ‘rbioacc’ provides ready-to-use functions to visualize and fully analyze such data. Under a Bayesian framework, this package fits a generic one-compartment toxicokinetic (TK) model automatically built from the data. It provides TK parameter estimates (appropriate uptake and elimination rates) and bioaccumulation metrics (e.g., BCF, BSAF, BMF). All parameter estimates, bioaccumulation metrics as well as predictions of internal concentrations into organisms are delivered with their uncertainty.
3. Bioaccumulation metrics are primarily provided in support of environmental risk assessment, in full compliance with regulatory requirements required to approve marketing applications of chemical substances.
4. This paper gives brief worked examples of the use of ‘rbioacc’ from data collected through standard bioaccumulation tests, and publicly available within the scientific literature. These examples constitute step-by-step user-guides to analyze any new data set, uploaded in the right format.

## Introduction

At least in part, Environmental Risk Assessment (ERA) is based on OECD regulatory guidelines to perform bioaccumulation tests (*e.g*., the test n°305 (Organisation for Economic Co-operation and Development, 2012) or the test n°315 (Organisation for Economic Co-operation and Development, 2008)). These documents specify the workflow to follow to obtain the bioaccumulation metrics on which regulatory decisions will be based. Indeed, bioaccumulation metrics are key criteria to define the capacity of chemicals to be bioaccumulated within organisms. This workflow can be performed with ‘rbioacc’, or directly on-line with the MOSAIC web platform at https://mosaic.univ-lyon1.fr/bioacc, making it easy for stakeholders to achieve this workflow. ‘rbioacc’ provides ready-to-use functions to visualize and fully analyze bioaccumulation test data. Such data are internal concentration regularly measured over time all along the experiment during which organisms are exposed to a chemical (exposure phase with accumulation) and then put into a clean media (depuration phase). ‘rbioacc’ automatically builds a generic one-compartment toxicokinetic (TK) model according to the input data, and provides TK parameter estimates (appropriate uptake and elimination rates) and bioaccumulation metrics (e.g., the Bio-Concentration Factor, or BCF) by fitting the TK model on data under a Bayesian framework. All parameter estimates and bioaccumulation metrics as well as predictions of internal concentrations in organisms are delivered with the quantification of their uncertainty. The overall uncertainty on TK parameter estimates is estimated as a joint posterior probability distribution, and marginally summarized for outputs as a median and a 95% credible interval (namely, the uncertainty range).

### Statement of need

‘rbioacc’ is an ‘R’-package compatible with ‘R’ version > 4.1.0 (R Core Team, 2021) and with all standard operating systems (macOS, Linux, Windows). All ‘rbioacc’ outputs have been compared with previously published results considering other TK model implementations under different software platforms. Giving very similar results, ‘rbioacc’ was thus confirmed as fit-for-purpose to fit TK models on bioaccumulation test data (Charles, Ratier, & Lopes, 2021; Charles, Ratier, Baudrot, et al., 2021; Ratier & Charles, 2021; Ratier, Lopes, Geffard, & Babut, 2021). All functions in ‘rbioacc’ can be used without a deep knowledge neither of the underlying probabilistic modelling or of the inference methodology. Indeed, they were designed to behave easily as possible, without requiring the user to provide prior values for input parameters. Meanwhile, models implemented in ‘rbioacc’ can be used as a first step to explore new models which could appear as more appropriate for some situations. Note that ‘rbioacc’ benefits from a user-friendly and freely available web interface, MOSAIC_bioacc_, from which the same analyzes can be reproduced directly on-line without having to implement them in ‘R’ programming. MOSAIC_bioacc_ is directly accessible from the MOSAIC platform at https://mosaic.univ-lyon1.fr (Charles, Veber, & Delignette-Muller, 2018) or directly from https://mosaic.univ-lyon1.fr/bioacc (Ratier, Lopes, Multari, et al., 2021).

### Availability

‘rbioacc’ is available as an official ‘R’-package directly downloadable from CRAN at https://CRAN.R-project.org/package=rbioacc. Other package dependencies and system requirements are documented to help users with their installation issues. Besides, ‘rbioacc’ is available on the GitHub repository https://github.com/aursiber/rbioacc/, especially for advanced users encountering troubles or having suggestions regarding the features or the hidden content of the ‘rbioacc’ functions.

### Installation

‘rbioacc’ is linked to Stan, a Bayesian sampler used to perform Bayesian inference with already implemented models, methods, and algorithms (Carpenter et al., 2017). Stan is fully integrated to ‘R’ via the ‘rstan’ (Stan Development Team, 2021) and ‘rstantools’ (Gabry, Goodrich, & Lysy, 2020) packages. In additional, to use ‘rbioacc’, you need to install all the other ‘R’-packages dependencies: ‘ggplot2’, ‘Rcpp’ (>= 0.12.0), ‘RcppParallel’ (>= 5.0.1), ‘testthat, ‘ggmcmc’, ‘GGally’, ‘loo’, ‘stringr’, ‘zoo’ (>= 1.8-9), ‘BH’ (>= 1.66.0), ‘RcppEigen’ (>= 0.3.3.3.0) ‘StanHeaders’ (>= 2.18.0)’, ‘knitr’, and ‘rmarkdown’. For this purpose, you can use the classical ‘R’ command that is provided in the ‘R’-script (Supplementary Information, SI). ‘rbioacc’ is also linked to the programming language C++ that speeds up calculations and runs simulations delivering the TK predictions. You should not have issues with C++ requirements since it is very well integrated in ‘R’.

### Main features

‘rbioacc’ allows to fit more complex TK models than those classically used today, offering the possibility to account for multiple exposure routes (up to four, among water, sediment, pore water and/or food) as well as the possibility of biotransformation of the parent chemical into several phase I metabolites and the potential dilution by growth of organisms (when growth measurements are available). The main functions in ‘rbioacc’ are ‘modelData()’ and ‘modelData_ode()’ to format and visualize raw data as well as to build the corresponding TK model. The ‘fitTK()’ function allows to fit a generic one-compartment TK model on the raw data; it provides estimates of the TK model parameters and the subsequent bioaccumulation metrics. Fitting outputs can be either displayed with ‘plot()’ or summarized with ‘quantile_table()’. For the calculation of bioaccumulation metrics, both the ‘bioacc_metric()’ and ‘plot()’ functions may be run delivering the plot of the posterior probability distribution of the bioaccumulation metric density. The ‘quantile()’ function gives a summary of this distribution with the median (50% quantile) and the uncertainty range (2.5% and 97.5% quantiles). The time at which 95% of the chemical is eliminated is provided by the ‘t95()’ function. Underlying equations of the fitted TK model, automatically built by the package from the input data, can be displayed and stored with the ‘equations()’ function, once the fitting process is complete. Other functions are available to check the goodness-of-fit (GoF) based on several criteria that we chose for their relevance and wide use in the Bayesian community: the Posterior Predictive Check (PPC, with the ‘ppc()’ function); the comparison between marginal prior and posterior probability distributions of the TK parameters (with the ‘plot_PriorPost()’ function); the correlation matrix between parameter estimates obtained from the joint posterior probability distribution (with the ‘corrMatrix()’ function); the correlation plot between parameter estimates displayed from the join posterior probability distribution (with the ‘corrPlot()’ function); the Potential Scale Reduction Factors of the parameters (PSRF, with the ‘psrf()’ function, expected to be < 1.01); the Widely Applicable Information Criterion (WAIC, with the ‘waic()’ function) for model comparisons on a same data set; and the Monte Carlo Markov Chain (MCMC) chains (n = 3 by default) with the ‘mcmcTraces()’ function. Additionally, the ‘predict()’ and ‘predict_manual()’ functions allow to perform predictions of internal concentrations within organisms, with or without previously observed data.

‘rbioacc’ currently handles constant and time-variable exposure concentration data according to a generic workflow:

1. Upload and format a data set;
2. Plot the data set;
3. Fit the appropriate TK model on the data;
4. Get model equations, parameter estimates and bioaccumulation metrics (chosen according to the exposure as automatically detected from the input data);
5. Get model predictions and their uncertainty superimposed to the data (Visual Posterior Check or VPC);
6. Check GoF criteria, prioritizing the PPC and the associated percentage of the observed data that are comprised within the 95% uncertainty range of their predictions (expected to be close to 95%);
7. Perform validation of the model comparing simulations under a new exposure concentration and externally collected data.

Those steps are described in full details in a ‘Tutorial’ available at https://doi.org/10.5281/zenodo.5092316, including the description on how to use all ‘rbioacc’ features. More information on the model and the inference process used in ‘rbioacc’ are given in an on-line ‘User Guide’ available at http://lbbe-shiny.univ-lyon1.fr/mosaic-bioacc/data/user_guide.pdf but also in our companion paper of the MOSAIC_bioacc_ web interface (Ratier et al., 2021). Please refer to this documentation for further introduction to the use of ‘rbioacc’.

The data, however, must be properly formatted beforehand, in order to be uploaded in the ‘R’ software (R Core Team, 2021) as a data frame from a .txt or a .csv file (that is comma, semicolon, or tabular separator). Each line of the data frame must correspond to a unique time point for a given replicate and a given exposure concentration of the contaminant. The data frame must contain at least four columns with predefined headings (Table 1): ‘time’, the time point of the measurement; ‘expw’, ‘exps’, ‘expf’, ‘exppw’: the exposure concentration, from water, sediment, food, and pore water, respectively; ‘replicate’, a unique integer or string for each replicate; ‘conc’, the measured internal concentrations. Further columns can exist: ‘concm*ℓ*’, for internal concentration measurements of metabolite *ℓ* (namely, ‘concm1’, ‘concm2’, …). Please note that only metabolites of phase I (deriving directly from the parent compound) in the metabolization process are considered; ‘growth’, for growth measurements (e.g., weight or size) of the organisms. Finally units must be fixed for ‘time’(in hours, minutes, days or weeks), for exposure concentrations (in *μg.mL*^−1^ or *μg.g*^−1^), for the measured concentrations (in *μg.g*^−1^), and for growth measurements (in g, mg, cm, mm or ‘other’). Replicate marking is dimensionless.

**Table 1.**
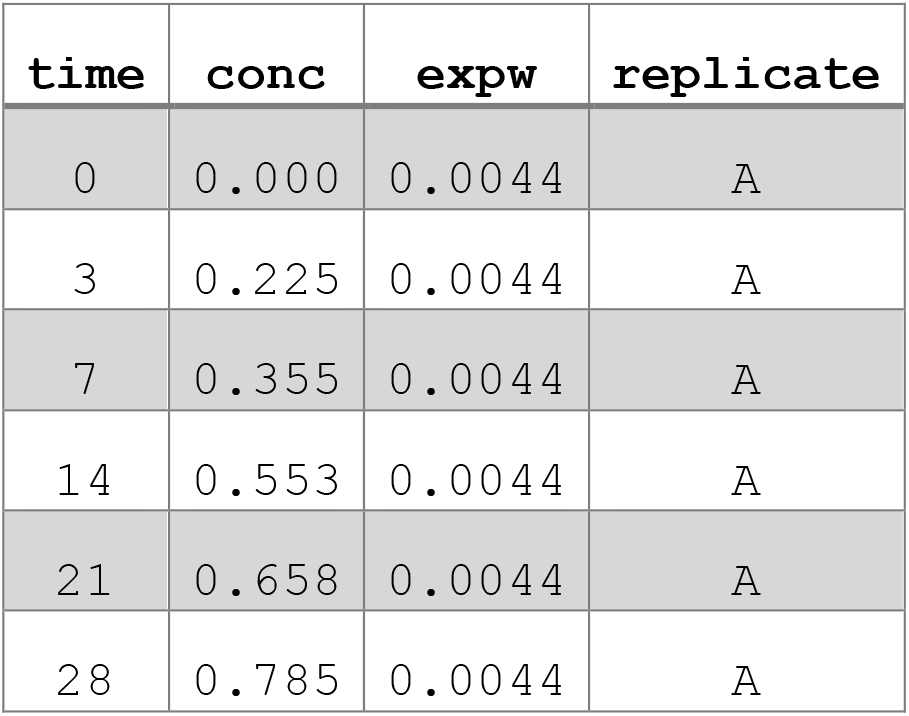
Example of a data set ready to be uploaded in ‘rbioacc’.

## Brief worked examples

### The ERA workflow

The ERA workflow proposed by the EFSA in the case of toxicokinetic-toxicodynamic (TKTD) models for the regulatory risk assessment of pesticides for aquatic organisms (Ockleford et al., 2018) consists of three steps: calibration, validation, and prediction. This workflow also applies to any TK model. The calibration step consists in fitting a TK model on bioaccumulation test data to get TK parameter estimates and deduce the bioaccumulation metrics: the BCF, the bio-sediment accumulation factor (BSAF) or the bio-magnification factor (BMF) depending on whether the exposure is by water, sediment, or food, respectively. All GoF criteria must be checked before proceeding to the validation step. The validation step consists in simulating the internal concentration over time as induced by the exposure to a chemical and comparing the predictions to newly collected bioaccumulation data. Results are validated from a VPC and 3 additional quantitative criteria (Ockleford et al., 2018). Finally, the prediction step consists in making simulations under more realistic exposure scenarios to assess the risk in closer relationship with the real world. For this last step which completes the entire workflow, it is of the utmost importance for the TK model to have been thoroughly calibrated and validated beforehand for the chemical-species combination under consideration.

To be in full compliance with regulatory guidelines and the above-mentioned workflow, the *modus operandi* with ‘rbioacc’ is illustrated below to be followed step-by-step with any other data set.

### Calibration step

To first illustrate the calibration step, we built two basic examples using standard bioaccumulation data sets available within ‘rbioacc’ thanks to the ‘data()’ function. The first data set was collected from a laboratory bioaccumulation test with the rainbow trout (*Oncorhynchus mykiss*) exposed to 2 different concentrations of a highly hydrophobic compound in spiked water for 49 days. There was 1 replicate per concentration. The duration of the depuration phase was 97 days. Internal concentrations were monitored at regular time points (Crookes & Brooke, 2011). The second example concerned the species *Chironomus tentans*, a freshwater invertebrate, exposed to benzo[a]pyrene in spiked sediment for 3 days. Only 1 exposure concentration was tested with 2 replicates. The duration of the depuration phase was 3 days. The internal concentrations for both the parent compound and its phase I metabolite were monitored at regular time points (Schuler, Wheeler, Bailer, & Lydy, 2003).

#### Upload data and fit the TK model accordingly

For the first example, because 2 different exposure concentrations were tested, one of them must be chosen before performing the analysis with ‘rbioacc’. For the second example, only 1 concentration has been tested; thus, the TK analysis can directly be launched with ‘rbioacc’. See the ‘R’-code given in SI to be copied and pasted, to identically reproduce all the results.

#### Results

The major results provided by ‘rbioacc’ are the predictions of the internal concentrations from the fitted model, superimposed to the observed data over time with the ‘plot()’ function. The posterior probability distribution of the bioaccumulation metric(s) is (are) given by the ‘bioacc_metric()’ function; this function applies whatever the exposure route(s) and the elimination process(es). If ‘bioacc_metric()’ is used with its default option, it provides the kinetic bioaccumulation metric(s). The steady-state bioaccumulation metric(s) is also available if the data show having more or less reached the steady-state at the end of the accumulation phase. A summary of the TK model parameter probability distributions is given by the ‘quantile_table()’ function. In addition, ‘rbioacc’ provides the equations of the underlying model, automatically built from the input data with the ‘equations()’ function. Figure 1 below shows the internal concentrations predicted from the fitted model and superimposed to the observed data for example 1: *Oncorhynchus mykiss* exposed to a hydrophobic chemical.

**Figure 1:**
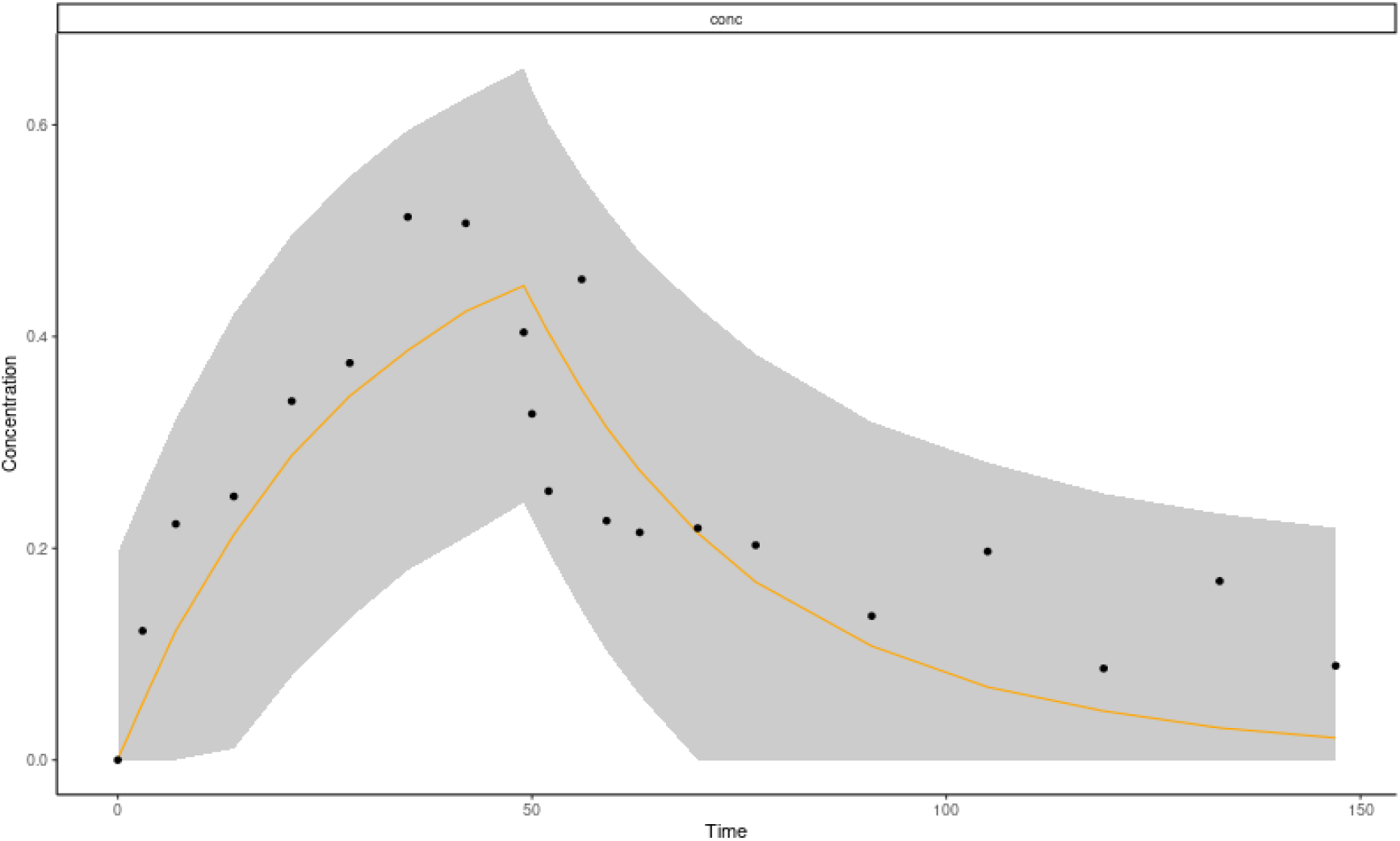
Fitting plot provided by ‘rbioacc’ for the example 1 with Oncorhynchus mykiss. Black dots are the observed data. The predicted median is symbolized by the orange plain line and the 95% uncertainty range by the grey area.

Figure 2 below shows the internal concentrations predicted from the fitted model and superimposed to the observed data for example 2: *Chironomus tentans* exposed to benzo[a]pyrene.

**Figure 2:**
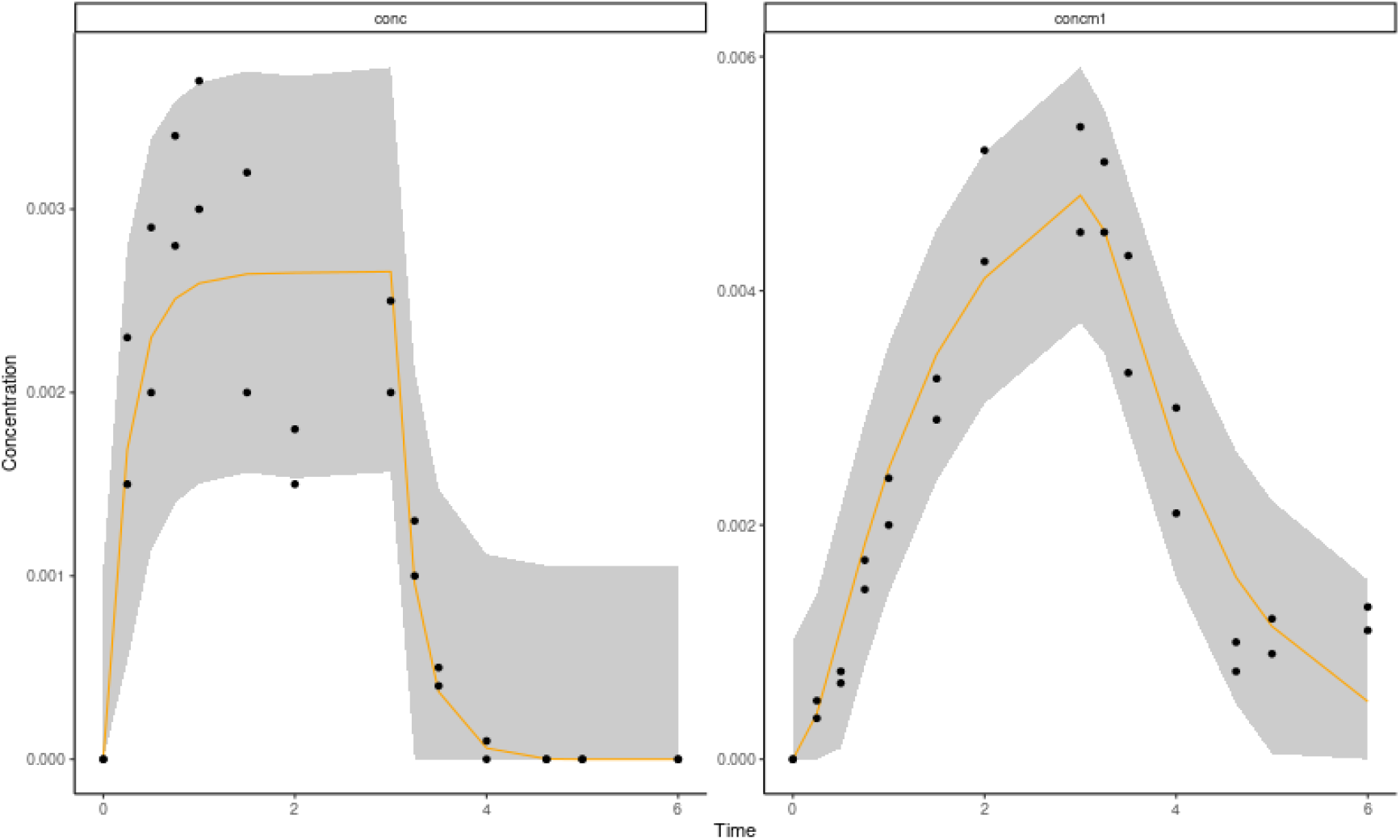
Fitting plots provided by ‘rbioacc’ for the example 2 with Chironomus tentans. Both fitting plots are displayed, for the parent compound (left) and its phase I metabolite (right). Black dots are the observed data. The predicted median curves are represented by the orange plain lines and the associated 95% uncertainty range by the grey areas.

#### Other goodness-of-fit criteria

Once the predictions of the internal concentrations are visually checked (VPC, Figs 1 & 2) several other GoF criteria require to be checked to evaluate how relevant are the fitting outputs, including the parameter estimates especially. For example, the PPC plot allows to compare each observed value to its prediction from the fitted TK model associated with its 95% uncertainty range. If the fitting process is correct, it is expected to have 95% of the observed values inside their 95% uncertainty range. With ‘rbioacc’, the PPC can be obtained with the ‘ppc()’ function as illustrated in Figure 3 below for example 2: we got 94.6% of data included within their uncertainty range. Additional GoF can also be checked, such as the comparison of prior and posterior probability distributions with the ‘plot_PriorPost()’ function, the correlation matrix of the model parameters (with the ‘corrMatrix()’ and/or ‘corrPlot()’ functions), the PSRF for each parameter with the ‘psrf()’ function, and the traces of the MCMC chains for each parameter with the ‘mcmcTraces()’ function (see SI). If several models have been fitted to the same data set then the WAIC can be asked for their comparison with the ‘waic()’ function.

**Figure 3:**
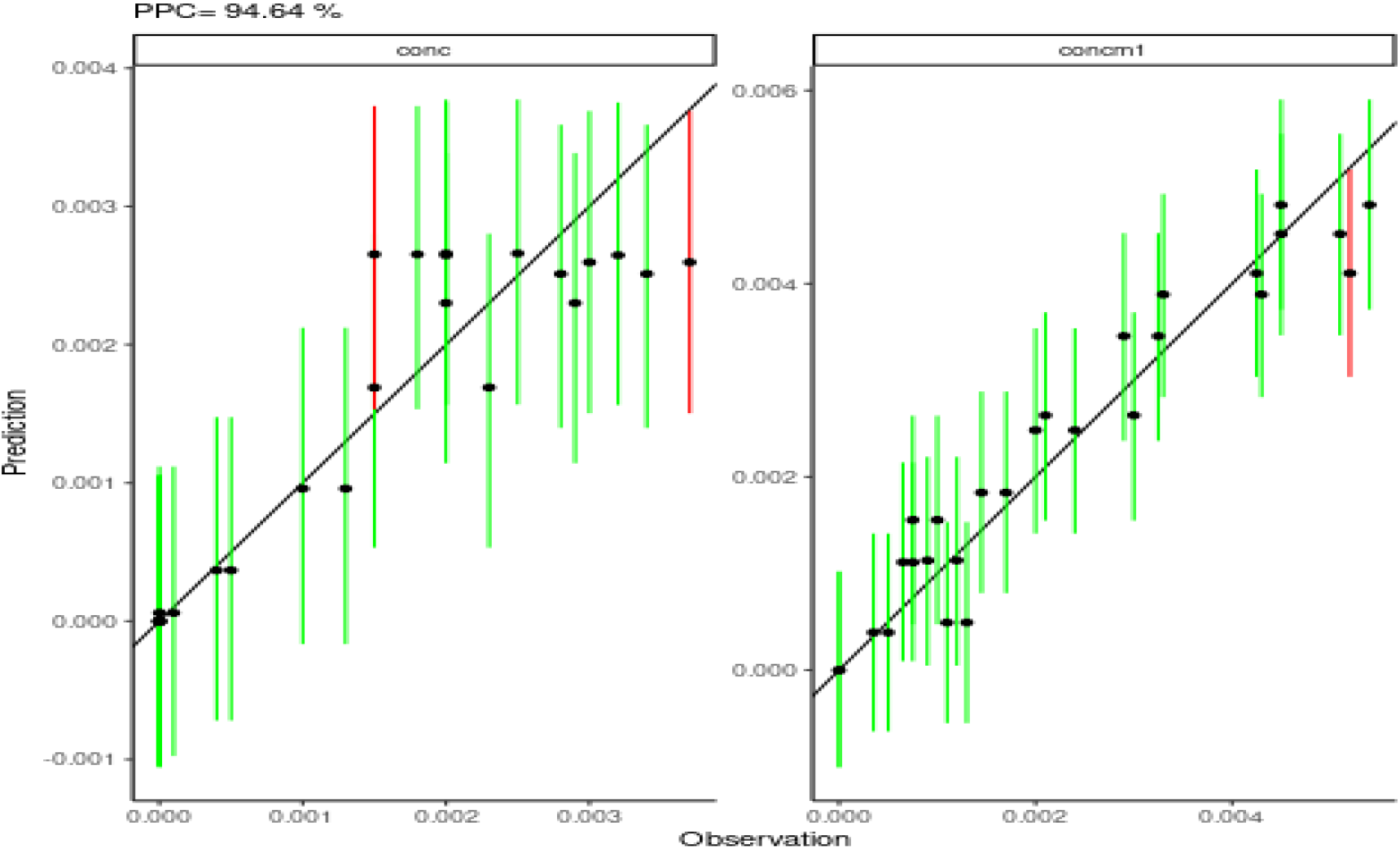
Posterior Predictive Check (PPC) as given by ‘rbioacc’ to check the goodness-of-fit for the example 2 with Chironomus tentans: PPC for the parent compound (left) and PPC for its phase I metabolite (right).

### Validation step

To illustrate the validation step with ‘rbioacc’, we chose a data set concerning *Spirostomum ambiguum* exposed to the pharmaceutical product fluoxetine at 0.025 *μg.mL*^−1^ under spiked water for 6 days (Nalecz-Jaweski, Wawryniuk, Giebultowicz, Olkowski, & Drobniewska, 2020). These data were first used to calibrate the appropriate TK model and to get the joint posterior probability distribution of the model parameters (results not shown). Once calibrated, the model was used to predict what may happen at 0.1 *μg.mL*^−1^ of exposure for which data were also available. Then, predictions were superimposed to these new data (Figure 4).

**Figure 4:**
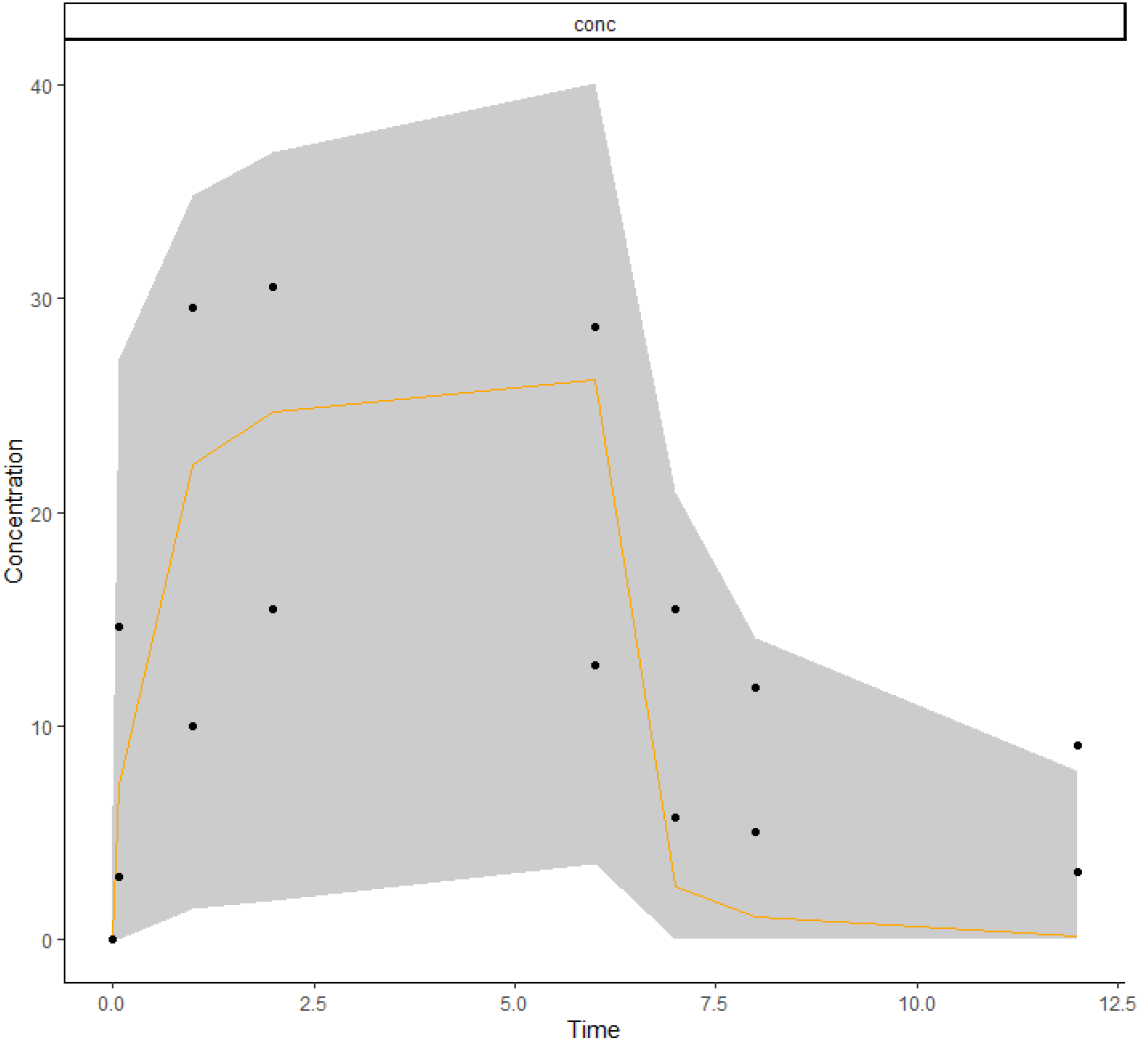
Example of a validation step for the example with Spirostomum ambiguum exposed to fluoxetine at 0.1 *μg.mL*^−1^. The orange curve is the predicted median of the internal concentration from the TK parameters estimated at 0.025 *μg.mL*^−1^ during a previous calibration step. The grey area is the 95% uncertainty range of the predictions, while black dots are the new observed data at 0.1 *μg.mL*^−1^.

### Prediction step

If the previous validation step provides relevant outputs, then the prediction step can be undertaken for further simulations at untested exposure concentrations or for a different accumulation time. In this perspective, both the ‘predict()’ or ‘predict_manual()’ functions of ‘rbioacc’ can be used. From the previous validation step with *Spirostomum ambiguum*, we performed simulation at 0.05 *μg.mL*^−1^ (Figure 5). This prediction was obtained by propagating the uncertainty on TK parameter estimates from the pre-exitent calibration step with *Spirostomum ambiguum*.

**Figure 5:**
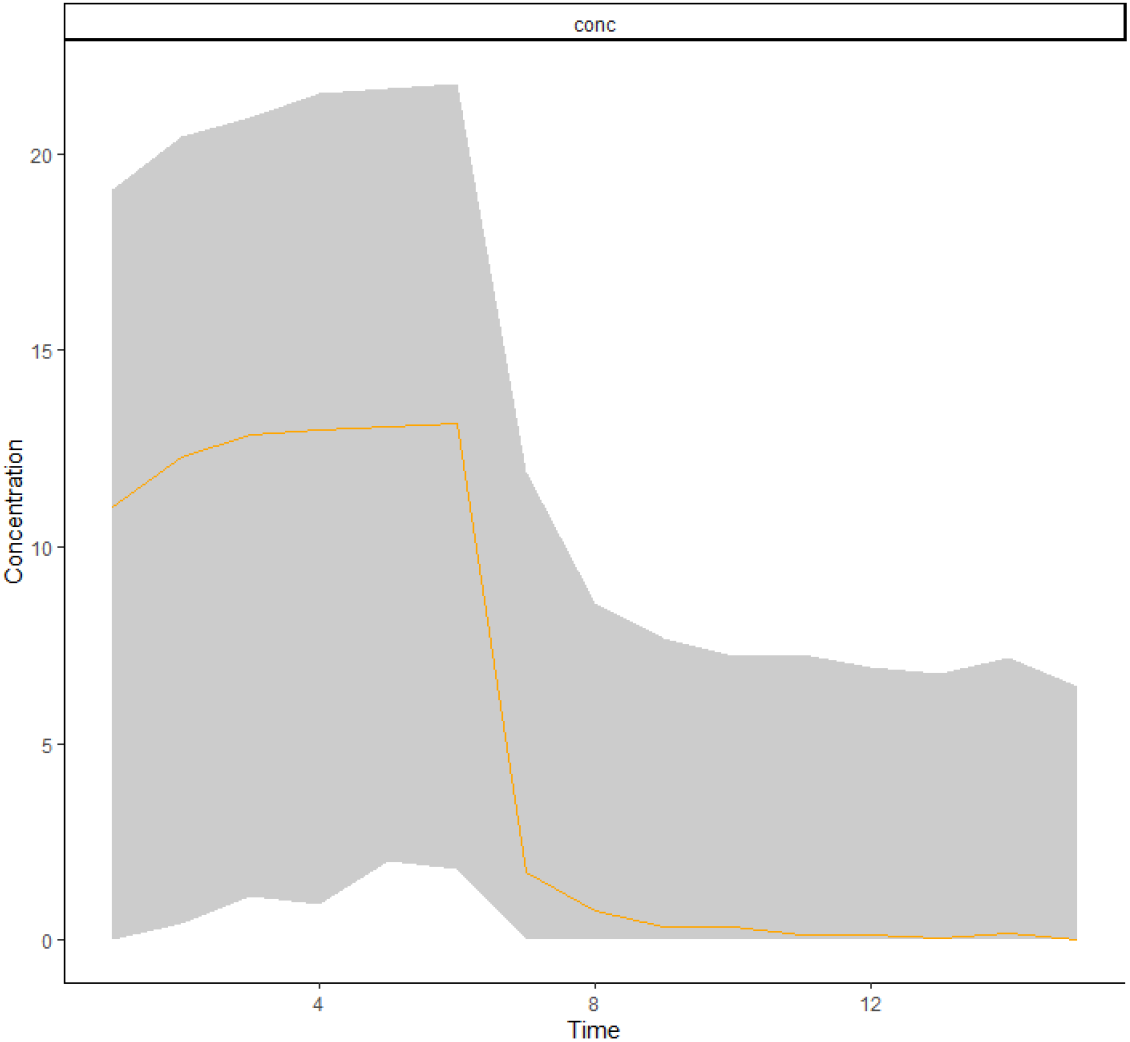
Example of a prediction step for the example with Spirostomum ambiguum exposed to fluoxetine at 0.05 *μg.mL*^−1^. The orange curve is the predicted median of the internal concentration with TK parameters estimated from data at 0.025 *μg.mL*^−1^. The gray area is the 95% uncertainty range around the median curve.

Another situation may require the use of the prediction step. Indeed, it may happen that only mean or median values for each parameter are available. These values may come from the scientific literature or from previous experiments for which the associated raw TK data have not been recorded. In such situations, neither calibration nor validation can be performed. Thus, only the ‘predict_manual()’ function can be used as illustrated below (Figure 6) from median values of TK parameters provided for *Oncorhynchus mykiss* by the authors themselves (Crookes & Brooke, 2011) at 0.01 *μg.mL*^−1^. The exposure concentration 0.01 *μg.mL*^−1^ was used in Figure 6, but the user can choose any other value, for example to design a new experiment for the same species-chemical combination. The counterpart of this approach is that no uncertainty can be propagated to the predictions (Figure 6).

**Figure 6:**
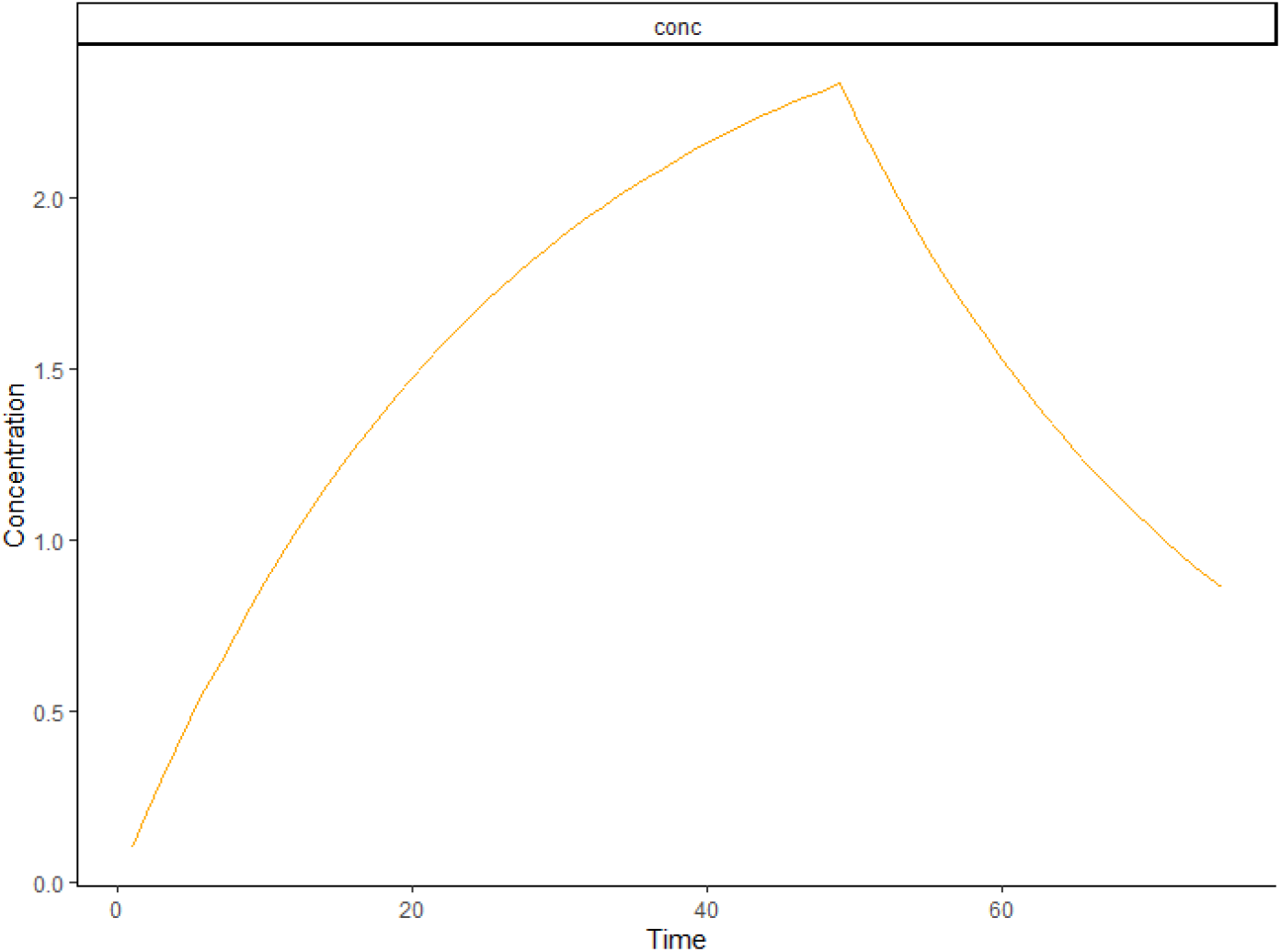
Example of a prediction step for fish Oncorhynchus mykiss exposed to a hydrophobic chemical. The orange curve is the predicted median curve of the internal concentration simulated from the median TK parameter values provided in the original paper at 0.01 *μg.mL*^−1^ (Crookes & Brooke, 2011).

## Bioaccumulation under time-variable exposure profiles

To mimic realistic environmental conditions, most often fluctuating than constant (*e.g*., when collecting field data), it may need to predict the internal concentration under time-variable exposure profiles (Ashauer & Brown, 2013; Baudrot & Charles, 2019). Such situations can easily be explored with ‘rbioacc’. First, it requires to upload two data files (one is expected when the exposure concentration is constant): one text file with the time-variable exposure concentrations over time (at least two columns entitled ‘time’ and ‘Cwater’; please notice that, despite this column heading, any exposure media can be considered, such as sediment, food or pore water) and another one with the corresponding internal concentration values over time (at least two columns entitled ‘time’ and ‘Cinternal’); these internal concentrations may have been previously simulated with the fitted corresponding TK model by ‘rbioacc’. Secondly, it requires to call the ‘modelData_ode()’ function which automatically manages the numerical integration of the ordinary differential equations that make the model behind and run simulations. Then the other functions symbolized can be used as illustrated above to get results and GOF criteria. The ‘R’-code in SI gives an example with a data set concerning *Sialis lutaria* exposed to a time-variable exposure profile of chlorpyrifos spiked water for 2 days (Rubach et al., 2010).

## MOSAIC_bioacc_: a web service interfacing ‘rbioacc’

### Brief presentation

The MOSAIC_bioacc_ web service has been designed to allow user-friendly TK modelling of complex situations in the field of ecotoxicology (Ratier, Lopes, Multari, et al., 2021). It is developed under the ‘R-shiny’ environment (Chang et al., 2021), providing an interactive interface to ‘rbioacc’. MOSAIC_bioacc_ also exploits the generic one-compartment TK model (Charles, Ratier, Baudrot, et al., 2021), meaning that organisms are considered as whole and unique compartments. Nevertheless, MOSAIC_bioacc_ proposes a more complex TK model than what is classically used today, offering the possibility to account for multiple exposure routes (up to four, among water, sediment, pore water and/or food) as well as the possibility of biotransformation of the parent chemical into several phase I metabolites; and the potential dilution by growth of organisms (growth measurements are additionally required). MOSAIC_bioacc_ automatically builds the TK model appropriate to the uploaded data, benefiting from the ‘rbioacc’ functions running behind on a dedicated server. MOSAIC_bioacc_ delivers all the GoF criteria for the user to easily assess the quality of the fitting results. It is a tool really thought for the regulatory domain, but that can also be helpful for the academic research or education.

### Last updates

Based on the clear observation that it is difficult to find enough raw data sets to validate generic modelling approaches, especially for TK models, we have recently developed a database with a collection of more than 200 accumulation-depuration data sets. They were extracted from the scientific literature and are now freely available on-line at http://lbbe-shiny.univ-lyon1.fr/mosaic-bioacc/data/database/TK_database.html. These data sets are associated with several outputs: the text file with raw data under the expected ‘rbioacc’ input format; the full report with the fitting results and the GoF criteria as provided either by MOSAIC_bioacc_ or ‘rbioacc’; and a link to the original paper (Ratier & Charles, 2021). MOSAIC_bioacc_ recently shifted from an all-embarked web application to be a user interface of the newly developed ‘rbioacc’ package. It allows all users to bring MOSAIC_bioacc_ and its self-explanatory functions to their own device for an easy integration of its methodology to every workflow. Furthermore, new features were added: a validation and prediction tools with the recent functions implemented in ‘rbioacc’. The present section showcases these very last updates of MOSAIC_bioacc_ still improving its intuitive use for any user without any need to invest in the underlying technicalities.

#### Aesthetic

The previous version of MOSAIC_bioacc_ was based on a scroll of only one web page. The new version works with several tabs unlocked once the user performed the actions required from the previous tab, except for the prediction and validation tabs that are always available. Globally, the same visual elements were kept, but they got reorganised in a tab structure with 5 different levels (Figure 7): ‘Data upload’, ‘Model and parameters’, ‘Results’, ‘Downloads’, ‘Prediction tool’. The results of the fitting process were also compacted in a tab and column structure to better suit the new no-scroll policy. This tremendously reduced the length of the previously displayed unique web page. This revamp in a tab based interface, as well as other minor changes, were partly decided upon by analysing the behaviour of the potential users of MOSAIC_bioacc_.

**Figure 7:**
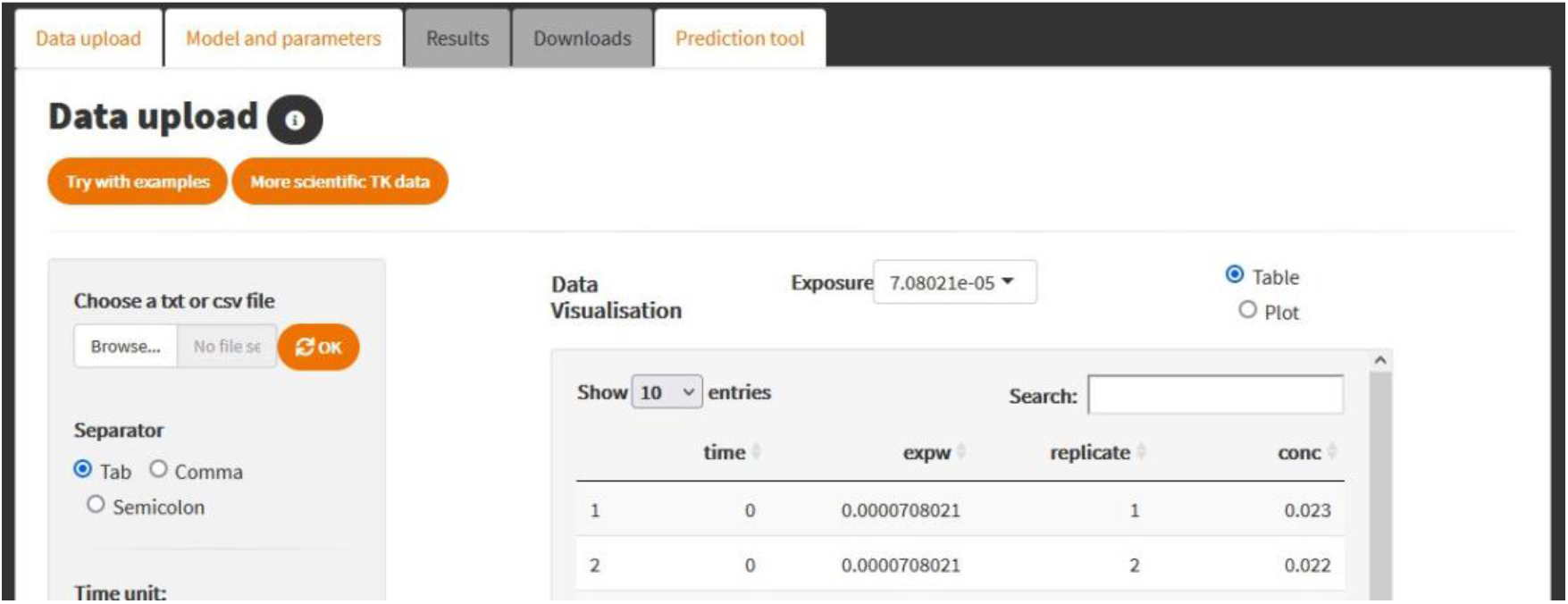
Visual homepage of the MOSAIC_bioacc_ first tab (‘Data upload’); once a data set has been uploaded, the second tab (‘Model and parameters’) is unlocked from which the TK model run can be asked thus unlocking ‘Results’ and ‘Downloads’ tabs in the following.

#### Architecture

The previous version of MOSAIC_bioacc_ suffered from a mix of classical and reactive ‘R’ coding, the biggest obstacle being the use of global variables. Global variables are generally not meant to be used in a reactive context, but can be forced if unavoidable for some very punctual cases. Using global variables allowed for a more classical way of coding, but it generated a series of scope related bugs, notably for the downloads, that needed to be fixed with lengthy patch ups. Furthermore, using global variables promotes excessive code re-evaluation, making the application less effective. Due to the long and iterative development process by different staff members, other issues appeared such as code duplication, repetitive operations on variables, creation of unused variables and non explicit, temporary variable names that ended up being kept. All in all, the ‘server’ file (‘server.R’, required by a ‘R-Shiny’ development) of the previous version of MOSAIC_bioacc_ was approximately 4.5k lines long. For all these reasons, this file was entirely rewritten, which allows an easier code to maintain, to understand and to upgrade, by sectioning the code in multiple files; by introducing documentation with the ‘roxygen2’ package (Wickham, Danenberg, Csardi, & Eugster, 2021) and more extensive comments through the code; and by strictly managing variable dependencies in an explicit way. With both the refactoring of the code and the integration symbolized, the ‘server.R’ file became approximately 1k lines long, 1.5k lines long by taking into account the other files containing code used by the server file itself. The major benefit to this restructuring was to include ‘rbioacc’ in the downloadable ‘R’-code from the application, gaining clarity and lines long, easier for a user to reuse the ‘R’ code.

#### New features

A validation and prediction tools were added to MOSAIC_bioacc_, for simulations of a chosen TK model with parameter values coming from either a previous calibration step, taken in the joint probability distribution of the estimated TK parameters, or manually entered by the user as point parameter values. So, the validation and the prediction tools allow the user to either propagate the uncertainties on parameters or not, in order to perform a simulation for a new exposure concentration. The validation tool takes things a step further by allowing the visual comparison of the predicted accumulation-depuration curve to newly observed validation data; in this case, the uncertainty on parameter estimates is required so that a previous calibration step needs to have been performed earlier.

## Conclusion

‘rbioacc’ was designed to help stakeholders and researchers to analyze TK data collected from bioaccumulation tests. This turnkey package automatically builds and fits the appropriate TK model, providing TK parameter estimates, GoF criteria and bioaccumulation metrics (namely, BCF/BSAF/BMF). In particular, it is designed to fulfil all requirements of regulators when examining marketing applications for chemical approval. In this perspective, the associated MOSAIC_bioacc_ web interface may reveals of particular help if supported by supervisory bodies.

## Declarations

## Acknowledgements

This work is part of the ANR project APPROve (ANR-18-CE34-0013) for an integrated approach to propose proteomics for biomonitoring: accumulation, fate and multi-markers (https://anr.fr/Projet-ANR-18-CE34-0013). This work benefited from the French GDR “Aquatic Ecotoxicology” framework which aims at fostering stimulating scientific discussions and collaborations for more integrative approaches. Authors fully thank all contributors to ‘rbioacc’: Théo Ciccia, Ophélia Gestin, Gauthier Multari and Alain Pavé. This work was performed using the computing facilities of the CC LBBE/PRABI.

## Author contributions

A.R.: supervision, formal analysis, data curation, writing manuscript. V.B. (main developer symbolized): conceptualisation, methodology, data curation, visualisation, writing manuscript. M.K.: data curation, conceptualisation, methodology, visualisation, writing manuscript. C.L.: supervision, formal analysis, data curation, reviewing manuscript. A.S.: conceptualisation, software maintenance, formal analysis, data curation, reviewing manuscript. S.C.: supervision, funding acquisition, project administration, formal analysis, data curation, reviewing manuscript.

## Supplementary information

All data sets used in this paper to illustrate the use of ‘rbioacc’ are available either within the package (with the ‘data()’ function), or within the public TK database at http://lbbe-shiny.univ-lyon1.fr/mosaic-bioacc/data/database/TK_database.html.

The ‘R’-script with all the ‘R’-commands allowing to exactly reproduce all the analyzes together with the resulting figures of the manuscript are available at our GitLab repository of the ‘rbioacc’ package: https://github.com/aursiber/rbioacc.

## References

Ashauer, R., & Brown, C. D. (2013). Highly time-variable exposure to chemicals--toward an assessment strategy. Integrated Environmental Assessment and Management, 9(3), e27–e33. doi:10.1002/ieam.1421

Baudrot, V., & Charles, S. (2019). Recommendations to address uncertainties in environmental risk assessment using toxicokinetics-toxicodynamics models. Scientific Reports, Natureresearch, 9, 11432. doi:http://dx.doi.org/10.1101/356469

Carpenter, B., Gelman, A., Hoffman, M. D., Lee, D., Goodrich, B., Betancourt, M., … Riddell, A. (2017). Stan: A probabilistic programming language. Journal of Statistical Software, 76(1), 1–32. doi:10.18637/jss.v076.i01

Chang, W., Cheng, J., Allaire, J., Sievert, C., Schloerke, B., Xie, Y., … Borges, B. (2021). ‘shiny’: Web Application Framework for ‘R’.

Charles, S., Ratier, A., Baudrot, V., Multari, G., Siberchicot, A., Wu, D., & Lopes, C. (2021). Taking full advantage of modelling to better assess environmental risk due to xenobiotics—the all-in-one facility MOSAIC. Environmental Science and Pollution Research. doi:10.1007/s11356-021-15042-7

Charles, S., Ratier, A., & Lopes, C. (2021). Generic solving of one-compartment toxicokinetic models. Journal of Exploratory Research in Pharmacology, submitted. doi:10.1101/2021.05.06.442956v1

Charles, S., Veber, P., & Delignette-Muller, M. L. (2018). MOSAIC: a web-interface for statistical analyses in ecotoxicology. Environmental Science and Pollution Research, 25, 11295–11302. doi:10.1007/s11356-017-9809-4

Crookes, M. J., & Brooke, D. N. (2011). Estimation of fish bioconcentration factor ( BCF ) from depuration data. Bristol, UK. Retrieved from https://www.google.com/url?sa=t&rct=j&q=&esrc=s&source=web&cd=&ved=2ahUKEwjioMzq1JPwAhUSyxoKHdFFDj8QFjAAegQIBBAD&url=https%3A%2F%2Fassets.publishing.service.gov.uk%2Fgovernment%2Fuploads%2Fsystem%2Fuploads%2Fattachment_data%2Ffile%2F291527%2Fscho0811buce-

Gabry, J., Goodrich, B., & Lysy, M. (2020). ‘rstantools’: Tools for Developing ‘R’ Packages Interfacing with ‘Stan’. Retrieved from https://cran.r-project.org/package=rstantools

Nalecz-Jaweski, G., Wawryniuk, M., Giebultowicz, J., Olkowski, A., & Drobniewska, A. (2020). Influence of Selected Antidepressants on the Ciliated Protozoan Spirostomum ambiguum: Toxicity, Bioaccumulation, and Biotransformation Products. Molecules, 25, 1476. doi:10.3390/molecules25071476

Ockleford, C., Adriaanse, P., Berny, P., Brock, T., Duquesne, S., Grilli, S., … EFSA PPR Panel. (2018). Scientific Opinion on the state of the art of Toxicokinetic/Toxicodynamic (TKTD) effect models for regulatory risk assessment of pesticides for aquatic organisms. EFSA Journal, 16(8), 5377. doi:10.2903/j.efsa.2018.5377

Organisation for Economic Co-operation and Development. (2008). Test No 315: Bioaccumulation in sediment-dwelling benthic oligochaetes. OECD Publishing. doi:10.1787/2074577x

Organisation for Economic Co-operation and Development. (2012). Test No. 305: Bioaccumulation in Fish: Aqueous and Dietary Exposure (Vol. 305). OECD Publishing. doi:10.1787/9789264185296-en

R Core Team. (2021). ‘R’: A Language and Environment for Statistical Computing. Vienna, Austria. Retrieved from https://www.r-project.org

Ratier, A., & Charles, S. (2021). Accumulation-depuration data collection in support of toxicokinetic modelling. Scientific Data, Nature, under mino(https://doi.org/10.1101/2021.04.15.439942). doi:10.1101/2021.04.15.439942

Ratier, A., Lopes, C., Geffard, O., & Babut, M. (2021). The added value of Bayesian inference for estimating biotransformation rates of organic contaminants in aquatic invertebrates. Aquatic Toxicology, 105811. doi:10.1016/j.aquatox.2021.105811

Ratier, A., Lopes, C., Multari, G., Mazerolles, V., Carpentier, P., & Charles, S. (2021). New perspectives on the calculation of bioaccumulation metrics for active substances in living organisms. Integrated Environmental Assessment and Management, on-line. doi:10.1101/2020.07.07.185835

Rubach, M. N., Ashauer, R., Maund, S. J., Baird, D. J., Van Den Brink, P. J., & den Brink, P. J. (2010). Toxicokinetic variation in 15 freshwater arthropod species exposed to the insecticide chlorpyrifos. Environmental Toxicology and Chemistry, 29(10), 2225–2234. doi:10.1002/etc.273

Schuler, L. J., Wheeler, M., Bailer, A. J., & Lydy, M. J. (2003). Toxicokinetics of sediment-sorbed benzo[a]pyrene and hexachlorobiphenyl using the freshwater invertebrates Hyalella azteca, Chironomus tentans, and Lumbriculus variegatus. Environmental Toxicology and Chemistry, 22(2), 439–449. doi:10.1897/1551-5028(2003)022<0439:TOSSBA>2.0.CO;2

Stan Development Team. (2021). ‘rstan’: the ‘R’ interface to ‘Stan’. Retrieved from https://cran.r-project.org/package=rstan

Wickham, H., Danenberg, P., Csardi, G., & Eugster, M. (2021). ‘roxygen2’’: In-Line Documentation for ‘R’.

